# Transcriptional read through interrupts boundary function in Drosophila

**DOI:** 10.1101/2023.02.16.528790

**Authors:** Olga Kyrchanova, Vladimir Sokolov, Maxim Tikhonov, Paul Schedl, Pavel Georgiev

**Affiliations:** Department of the Control of Genetic Processes, Institute of Gene Biology Russian Academy of Sciences, 34/5 Vavilov St., Moscow 119334, Russia; Center for Precision Genome Editing and Genetic Technologies for Biomedicine, Institute of Gene Biology, Russian Academy of Sciences, 34/5 Vavilov St., Moscow 119334, Russia; Department of Molecular Biology, Princeton University, Princeton, NJ, 08544, USA

**Keywords:** chromatin boundary, insulator, Bithorax complex, Abd-B, scs, Fub, Fab-7, transcription, lncRNA

## Abstract

In higher eukaryotes enhancer-promoter interactions are known to be restricted by the chromatin insulators/boundaries that delimit topologically associated domains (TADs); however, there are instances in which enhancer-promoter interactions span one or more boundary elements/TADs. At present, the mechanisms that enable cross-TAD regulatory interaction are not known. In the studies reported here we have taken advantage of the well characterized *Drosophila* Bithorax complex (BX-C) to study one potential mechanism for controlling boundary function and TAD organization. The regulatory domains of BX-C are flanked by boundaries which function to block crosstalk with their neighboring domains and also to support long distance interactions between the regulatory domains and their target gene. As many lncRNAs have been found in BX-C, we asked whether transcriptional readthrough can impact boundary function. For this purpose, we took advantage of two BX-C boundary replacement platforms, *Fab-7^attP50^* and *F2^attP^*, in which the *Fab-7* and *Fub* boundaries, respectively, are deleted and replaced with an *attP* site. We introduced boundary elements, promoters and polyadenylation signals arranged in different combinations and then assayed for boundary function. Our results show that transcriptional readthrough can interfere with boundary activity. Since lncRNAs represent a significant fraction of Pol II transcripts in multicellular eukaryotes, it is possible that many of them may function in the regulation of TAD organization.

**Author Summary:** Recent studies have shown that much genome in higher eukaryotes is transcribed into non-protein coding lncRNAs. It is though that lncRNAs may preform important regulatory functions, including the formation of protein complexes, organization of functional interactions between enhancers and promoters and the maintenance of open chromatin. Here we examined how transcription from promoters inserted into the *Drosophila* Bithorax complex can impact the boundaries that are responsible for establishing independent regulatory domains. Surprisingly, we found that even a relatively low level of transcriptional readthrough can impair boundary function. Transcription also affects the activity of enhancers located in BX-C regulatory domains. Taken together, our results raise the possibility that transcriptional readthrough may be a widely used mechanism to alter chromosome structure and regulate gene expression.

## Introduction

The chromosomes of multicellular animals are organized into a series of looped domains called TADs (topologically associated domains) [1–4]. While a variety of elements contribute to folding the chromatin fiber (e.g., the tethering elements that help link enhancers to promoters [5], this 3-dimensional organization depends, in part, on special elements called boundaries or insulators [1,6,7]. Although boundary elements have now been identified in many different species, they have been most thoroughly characterized in *Drosophila* [1,7,8]. Fly boundaries are 150 bp to 1.5 kb in length and span one or more nucleosome free nuclease hypersensitive regions that are formed by different combinations of chromosomal architectural proteins including *Drosophila* CTCF (dCTCF) [9,10].

Functional studies using transgene assays indicate that in addition to subdividing the chromosome into a series of looped domains, fly boundary elements have genetic functions [7,11,12]. When placed between an enhancer or silencer and a reporter, they prevent regulatory interactions. When a reporter is bracketed by boundary elements, they protect against chromosomal position effects. With some exceptions, boundary function in these assays is “constitutive” –i.e., it is observed throughout development and is independent of cell type. The likely reasons for this constitutive activity is that most of the fly architectural proteins are ubiquitously expressed [8] and that different combinations of these proteins are deployed to generate the activity of individual boundaries [13,14].

Since multiple functionally redundant architectural proteins contribute to the functions of individual fly boundaries in flies, it seems unlikely that TADs will undergo genome-wide reorganization during cellular differentiation as this would require a change in the patterns of expression of multiple chromosomal proteins. Rather, one might expect that TAD organization would be subject to local alterations by modulating the insulating functions of specific boundary elements. In the studies reported here we have used the *Drosophila* bithorax complex (BX-C) to identify mechanisms for modulating local boundary function.

BX-C is responsible for specifying the nine posterior-most parasegments (PS5-PS14 in embryo) (segments T3-A9 in adults) of fly [15–18]. As there are three homeotic genes in BX-C: *Ultrabithorax (Ubx), abdominal-A (abd-A)* and *Abdominal-B (Abd-B)* (Fig. 1A) parasegment (segment) specification depends up modulating their expression in patterns appropriate for the proper differentiation of each parasegment. This is accomplished by subdividing the complex into nine *cis*-regulatory domains. Each domain has tissue and stage specific enhancers responsible for directing a unique parasegment specific pattern of expression of one of the homeotic genes [16,19–23]. *Ubx* is responsible for specifying PS5(T3) and PS6 (A1) and its expression in these two parasegments is controlled by the *bx/abx* and *bxd/pbx* regulatory domains respectively. The *infra-abdominal* (*iab*) domains regulate the transcription of *abd-A* and *Abd-B*. The *abd-A* gene is controlled by *iab-2, iab-3*, and *iab-4* in PS7 (A2), PS8 (A3), and PS9 (A4), respectively. Four domains, *iab-5, iab-6, iab-7*, and *iab-8,9*, regulate *Abd-B* expression in PS10 (A5), PS11 (A6), PS12 (A7), and PS13,14 (A8 (♀), A9 (♂)), respectively (Fig. 1A). The regulatory domains are activated sequentially in successive parasegments along the anterior-posterior axis [17]. The activity state, *on* or *off*, of the regulatory domains is set early in development by maternal, gap and pair-rule gene proteins which bind to initiation elements in each domain [24–26]. Once the activity state is set it is remembered during the remainder of development by mechanisms that depend upon trithorax and Polycomb group proteins[27–30].

**Fig. 1.**
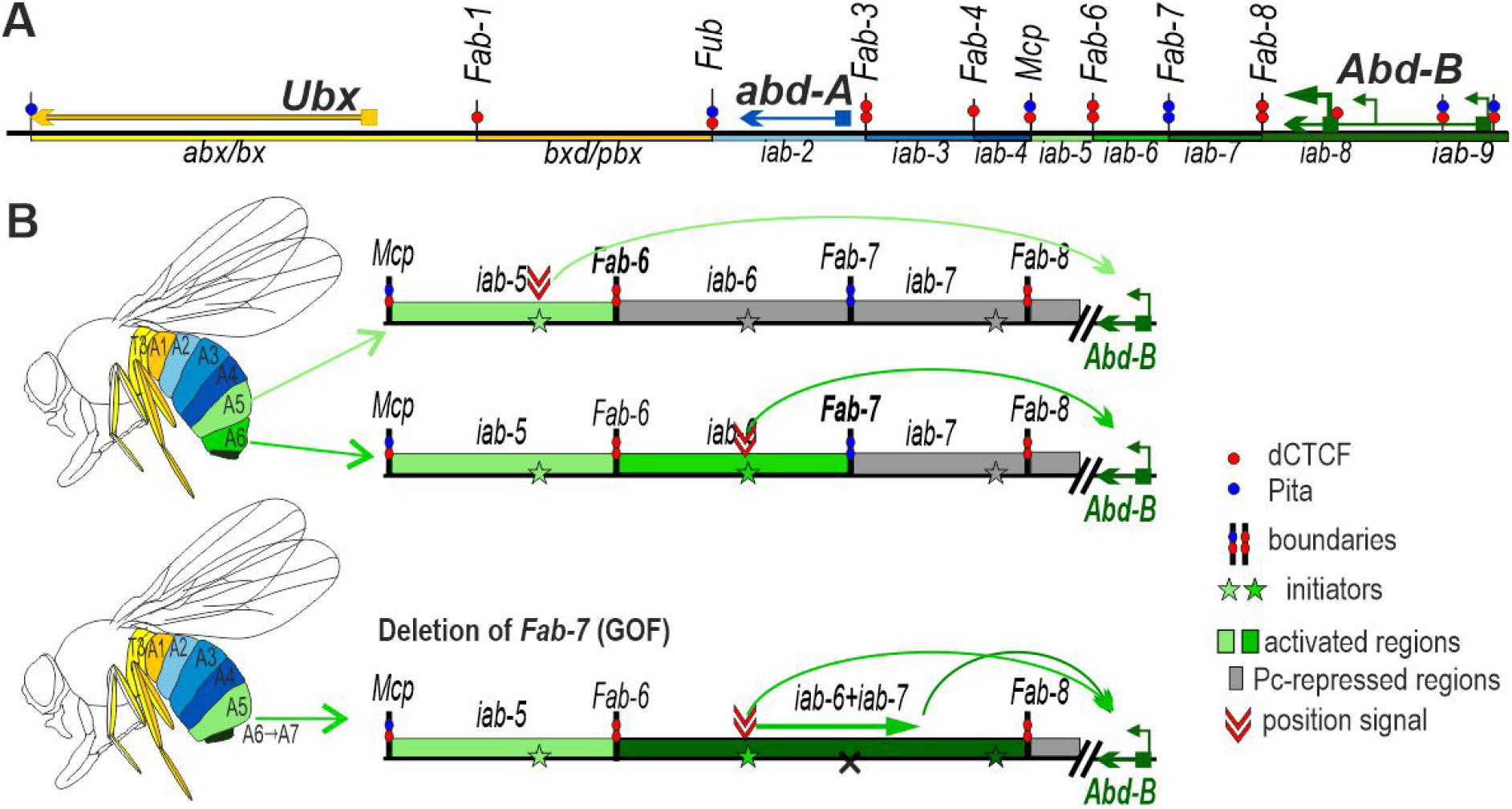
Boundaries organize enhancer-promoter interactions in the *Abd-B* gene of the BX-C. (A) Map of the BX-C showing the location of the three homeotic genes and the parasegment-specific regulatory domains. There are nine *cis*-regulatory domains (shown as colored boxes) that are responsible for the regulation of the BX-C genes and the specification of parsegments 5 to 13, which correspond to T3-A8 segments. The *abx/bx* (yellow) and *bxd/pbx* (orange) domains activate *Ubx*, *iab-2 – iab-4* (shades of blue) – *abd-A* and *iab-5–9* (shades of green) – *Abd-B*. Lines with colored circles mark chromatin boundaries. The dCTCF, Pita, and Su(Hw) binding sites at the boundaries are shown as red, blue, and yellow circles, respectively. (B) Schematic presentation of *Abd-B* activation in A5(PS11) and A6(PS12) segments (parasegments). (C) Deletion of the *Fab-7* boundary results in premature activation of the iab-7 domain in A6 (PS12).

Each domain is flanked by boundary elements which function to block crosstalk between initiation elements in adjacent regulatory domains [26,27,31–36]. For example, the *Fab-7* boundary in the *Abd-B* region of BX-C separates the *iab-6* and *iab-7* domains. When *Fab-7* is deleted *iab-6* and *iab-7* fuse into a single domain and the *iab-6* initiation element inappropriately activates *iab-7* in PS11 (A6) (Fig. 1B). As a result, *iab-7* drives *Abd-B* expression not only in PS12(A7) but also PS11(A6) transforming PS11(A6) into a copy of PS12 (A5). Most BX-C boundaries also have a second function, which is boundary “bypass”. For example, the *Abd-B* regulatory domains *iab-5, iab-6* and *iab-7* are separated from their gene target by one or more boundaries (Fig. 1A). In order for these domains to regulate *Abd-B* there must be a mechanism that enables the enhancers in each domain to bypass the intervening boundaries. Recent studies have shown that the *Fab-7* and *Fab-8* boundaries have elements that confer bypass activity enabling the domain immediately distal to the boundary to “jump over” the intervening boundaries and activate *Abd-B* expression [37–39].

In previous studies [40,41], the *Fab-7* boundary was replaced with two versions of the *scs* insulator from the 87A7 heat shock locus, *scs* and *scs^min^* [42–44]. The larger version, *scs*, is a complex insulator containing the *Cad87A* and *CG31211* promoters (Fig. 2A). The smaller fragment, *scs^min^*, lacks most of the *Cad87A* promoter [44] (Fig. 2A). Hogga et al. 2002 showed that when the *Cad87A* promoter in the larger *scs* replacement is oriented towards *iab-6* it disrupts the functioning of the *iab-6* and *iab-5* regulatory domains inducing a loss-of-function (LOF) phenotype in which the A6 and A5 cuticle had morphological features like A4 [40]. They suggested that readthrough transcription from the *Cad87*_promoter inactivates enhancers in *iab-5* and *iab-6* required for the development of the adult cuticle. However, a completely different result was observed when the *Cad87* promoter in the *scs* replacement was oriented towards *iab-7*. In this case, a gain-of-function (GOF) phenotype was induced: A6 was converted into a copy of A7. It was thought that transcription into *iab-7* disrupted Polycomb dependent silencing, but did not impact the activity of the *iab-7* tissue/stage specific enhancers. As these results seemed inconsistent, we decided to reinvestigate the functioning of both *scs* and *scs^min^* replacements in *Fab-7*. In the course of our studies, we discovered that transcriptional readthrough appears to be general mechanism for turning boundary function off, and thus is likely to have an important role in both remodeling TAD organization and change patterns of gene regulation.

**Fig. 2.**
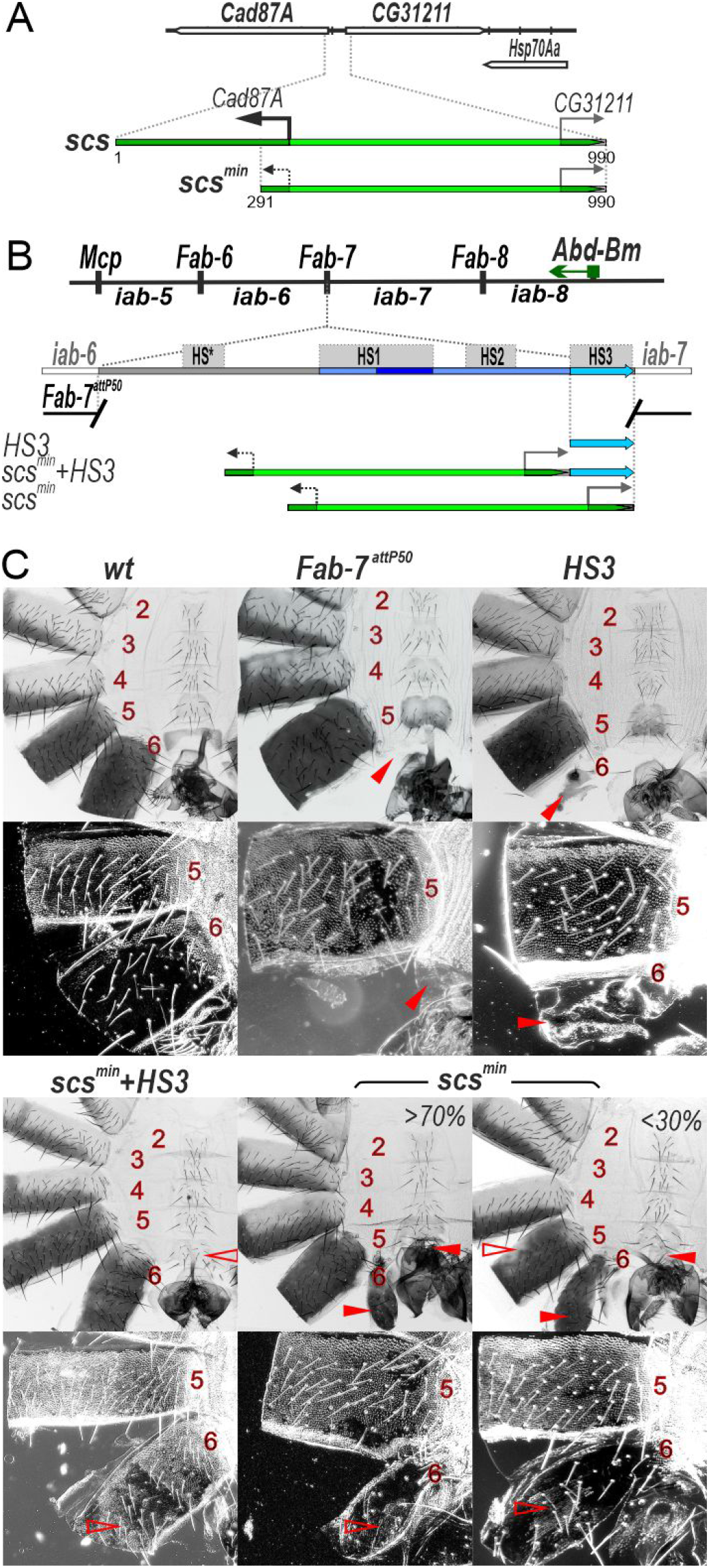
HS3 (*iab-7* PRE) is required for boundary activity of the *scs^min^* insulator. (A) Schematic presentation of the scs insulator (marked as green line). TSS - transcription start sites are marked by magenta arrows. PAS is designated as a STOP signal. Exons of *Cad87A* and *CG31211* are marked as white boxes with brown hatching. The black arrow indicates the *Cad87A* promoter that is active in embryos [40]. (B) Schematic representation of the *abd-A - Abd-B* regulatory regions and *Fab-7^attP50^* platform in which four hypersensitive sites, HS*, HS1, HS2, and HS3 (marked as grey boxes) are deleted. The endpoints of the deletion are indicated by breaks in the red line. HS3 marked as a blue line. Replacement fragments are shown below the map with a summary of their cuticle (tergite and sternite) phenotypes. (C) Bright field (top) and dark field (bottom) images of cuticles prepared from males of *wild type (wt), Fab-7^attP50^, HS3, scs^min^, scs^min^+HS3* transgenic lines. Abdominal segments are numbered. The filled red arrowheads show morphological features indicative of GOF transformations. The empty red arrowheads show the signs of the LOF transformation, which is directly correlated with boundary function.

## Results

### The *scs^min^* insulator can block crosstalk between the *iab-5* and *iab-6* domains only in cooperation with the *iab-7* PRE

In the boundary replacement experiments of Hogga et al [40,41] the *scs^min^* insulator was introduced into a *Fab-7* deletion which removes three of the four nuclease hypersensitive sites associated with the *Fab-7* boundary, HS*, HS1 and H2. This deletion results in an incomplete GOF transformation of A6 (PS11) into A7 (PS12). When *scs^min^* replaced this deletion, it blocked crosstalk between the *iab-6* and *iab-7* initiation elements and rescued the GOF phenotype of the starting deletion. However, since *scs^min^* does not support boundary bypass, the A6 segment was transformed towards A5.

HS3 was retained in the starting *Fab-7* deletion was found to induce Polycomb-dependent silencing and was thought to function as the *iab-7* Polycomb Response Element (*iab-7* PRE) [36,45,46]. However, recent studies showed that HS3 has insulating activity [47] and that this is likely its primary function. In fact, a fully functional *Fab-7* boundary can be generated by combining HS3 with the distal half of HS1 (dHS1) [37,47]. This finding made us wonder whether *scs^min^* would be able to block crosstalk between *iab-6* and *iab-7* in the absence of HS3.

To address this question, we used a previously characterized *Fab-7^attP50^* replacement platform in which the HS*, HS1, HS2 and HS3 are substituted by an *attP* site [48] (Fig. 2B). In the starting *Fab-7^attP50^* deletion *iab-7* is inappropriately activated in A6(PS11) and this results in the transformation of A6 into a copy of A7 so that both the A6 and A7 segments are absent in adult males (Fig. 2C).

To test for *scs^min^* function with and without HS3 we introduced three replacements, *scs^min^+HS3*, *HS3* and *scs^min^* into the *Fab-7^attP50^* platform. Though *scs^min^+HS3* has a slightly different sequence composition than the previously described *scs^min^* replacement [41] due to the use of different replacement platforms, its activity is similar. As shown in Fig. 2C, the *scs^min^+HS3* combination rescues the GOF transformation of A6 in adult males and the A6 segment is present. However, because this combination lacks bypass activity, the *iab-6* regulatory domain is blocked from regulating *Abd-B* expression in A6. As a consequence, the morphology of the A6 segment resembles that in A5. Instead of a banana shape without any bristles, the sternite has a quadrilateral shape and is covered in bristles and it resembles the sternite in A5. In *wild type* (*wt*) the trichome hairs on the A6 tergite are restricted to the anterior and ventral margin, while the trichome hairs cover most all of the A5 tergite (Fig. 2C). As can be seen in the darkfield image, the A6 tergite in *scs^min^+HS3* males is covered with trichome hairs just like A5. The same phenotypes were observed in patches of unpigmented cuticle. These transformations will be considered further below. As was observed for *Fab-7* (class II) deletions that retained HS3 [36], the *HS3* replacement alone has only a limited ability to block crosstalk between *iab-6* and *iab-7*. In *HS3* males, the A6 tergite is greatly reduced in size and the A6 sternite is completely missing [36,47] (Fig. 2C).

Like HS3, the *scs^min^* replacement only partially blocks crosstalk between *iab-6* and *iab-7* (Fig. 2C). However, it differs from the HS3 replacement in that there are a range of phenotypes in adult *scs^min^* males. In all *scs^min^* males there is a residual A6 tergite, while the A6 sternite is absent. The residual tergite has patches of cells with trichome hairs indicating that in these cells there is a LOF transformation in parasegment/segment identify from PS11/A6 to PS10/A5. In about 30% of the males, the morphogenesis of A5 is also affected. As shown in Fig. 2C, there are patches of tissue in the A5 tergite that are not fully pigment. This phenotype indicates that the *iab-5* regulatory domain is not fully functional in a subset of *scs^min^* males and we will return to this issue below.

### Transcription induced by the *Cad87A* promoter in *scs* can affect the activity of the *iab-7* domain

The finding that *scs^min^* must be combined with HS3 to efficiently block crosstalk between *iab-6* and *iab-7* prompted us to examine the blocking activity of the larger *scs* fragment which contains the *Cad87A* promoter (Fig. 3A). In previous study [40], when *scs* is inserted in the reverse orientation (*scs^R^*) so that the promoter is directed towards *iab-7*, males have a GOF phenotype in which A6 is transformed into A7. We repeated this experiment by inserting the *scs^R^+HS3* combination into *Fab-7^attP50^* and we observed similr GOF phenotypes (Fig. 3B). To explain the GOF transformation, it was suggested that transcription from the *Cad87A* promoter through *iab-7* domain induced the premature activation of *iab-7* by suppressing Polycomb (Pc)-dependent silencing [40]. However, since *scs^min^* cannot efficiently insulate *iab-6* from *iab-7* in the absence of HS3, an alternative possibility is that transcription through HS3 from the *Cad87A* promoter disrupts the boundary and/or PRE activity of HS3.

**Fig. 3.**
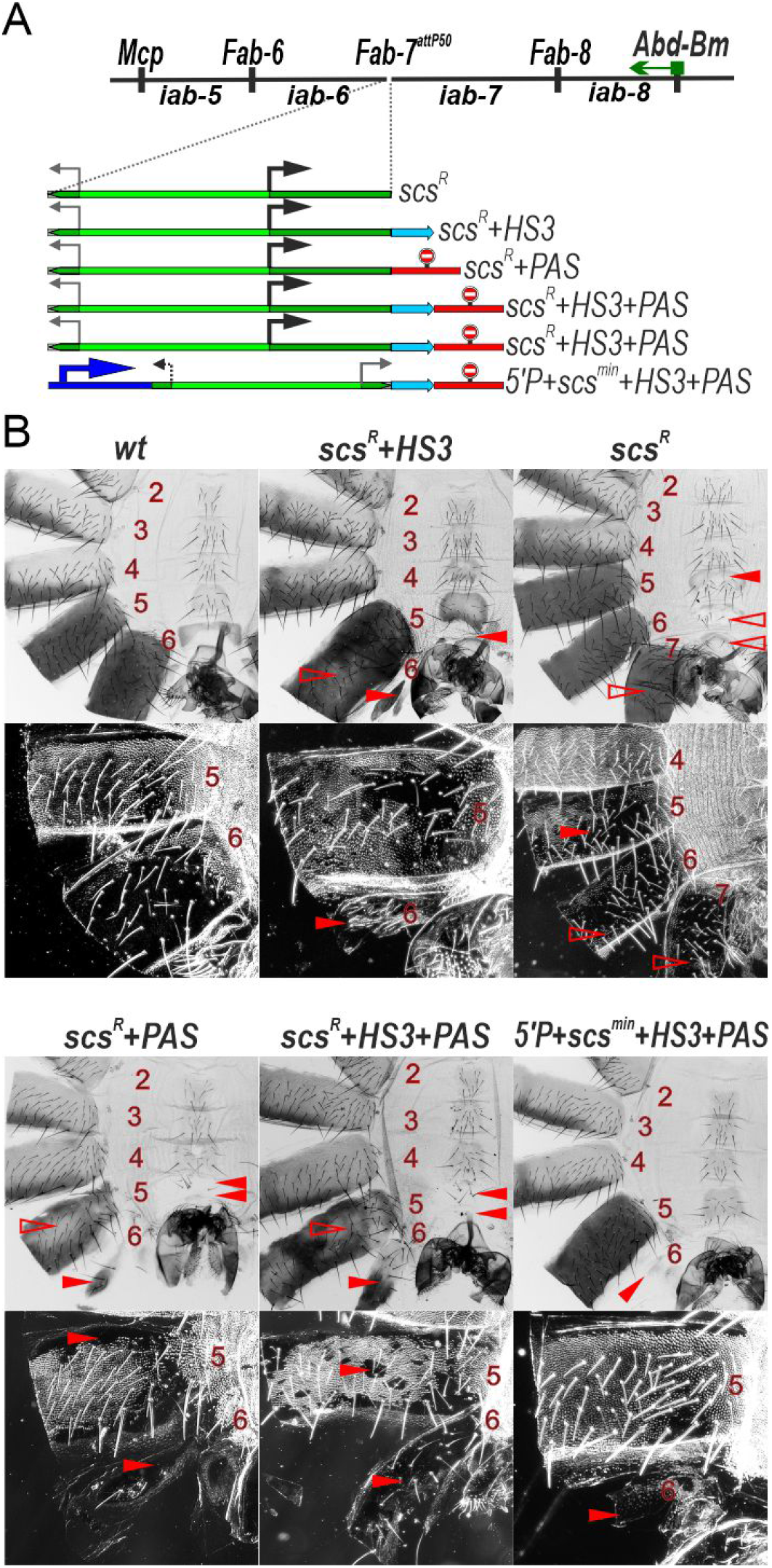
The *Cad87A* promoter in *scs* inserted in the reverse orientation (*scs^R^*) is responsible for inactivation of the *iab-7* enhancers. (A) *Fab-7* boundary replacement schemes in which *scs* was inserted in reverse orientation. (B) Bright field (top) and dark field (bottom) images of cuticles prepared from males of *wt*, *scs^R^+HS3, scs^R^, scs^R^+PAS, scs^R^+HS3+PAS, 5’P+scs^min^+HS3+PAS. (β 2 ряда – 6 линий*). Other designations are as in Fig 2.

To distinguish between these two models, we inserted *scs* in the reverse orientation in the *Fab-7^attP50^* platform. Fig. 3B shows that in the absence of HS3, *scs^R^* has a completely different phenotype. Instead of a GOF transformation of A6 into a copy of A7, there is an LOF transformation in which A7 is transformed towards A6. Unlike *wt* males, which lack an A7 segment, *scs^R^* males have an A7 tergite and sternite. The tergite is fully pigmented and the trichome hairs are largely (but not completely) restricted to the anterior and dorsal edges. The sternite has a banana shape like the sternite in A6; however, it is malformed and has bristles. These results suggest that transcription initiated from the *Cad87A* promoter leads to an inactivation of the *iab-7* enhancers. The phenotypic abnormalities of *scs^R^* are not restricted to A7. The A6 tergite has ectopic patches of trichome hairs, while the A6 sternite has bristles and is misshapen. While A6 shows evidence of a LOF transformation in portions of the adult cuticle, the opposite effect is observed in A5. The A5 tergite is partially devoid of trichome hairs, while A5 sternite has an abnormal banana like shape.

### Transcription disrupts boundary function

Taken together these findings suggest that the GOF transformations observed previously [40] in the *scs^R^* replacement might be due to the transcription induced inactivation of HS3 boundary activity rather than a transcription induce activation of *iab-7* in A6. This new model makes several predictions which we have tested. First, it should be possible to rescue the LOF phenotypes in A7 of the *scs^R^* replacement by introducing a transcription termination element, polyadenylation signal (*PAS*), in between *scs^R^* and the *iab-7* domain. If the blocking activity of the *scs^R^* element on its own is not too much different from *scs^min^*, then there should be a GOF transformation of A6 towards A7. This is what we observe in *scs^R^+PAS* males (Fig. 3B). There is a rudimentary A6 tergite with patches of ectopic trichome hairs, while the A6 sternite is absent.

Second, if transcription from the *Cad87* promoter in *scs* into *iab-7* results in a premature activation of the *iab-7* domain we should be able to block this activation by introducing the *PAS* element downstream of the *scs^R^+HS3* combination. In this case, the *scs^R^+HS3* combination would be expected to be functional (just like *scs^min^+HS3*) and block crosstalk between the *iab-6* and *iab-7* domains rescuing the GOF transformation of *Fab-7^attP^*. On the other hand, if transcriptional readthrough of *HS3* disrupts its ability to complement the *scs* element, then we should observe a GOF transformation of A6 toward A7 as was reported previously [40]. Fig. 3B shows that the later prediction is correct. Like *scs^R^+PAS*, male flies carrying the *scs^R^+HS3+PAS* combination have a GOF transformation of A6 towards A7 (Fig. 3B).

To further test the idea that transcriptional readthrough can disrupt boundary function, we generated a quadripartite replacement, *5’P+scs^min^+HS3+PAS*, consisting of the P-element promoter (*5’P*). As shown in Fig. 3B, inclusion of the P-element promoter disrupts the boundary activity of *scs^min^+HS3*. While *scs^min^+HS3* on its own rescues the GOF transformation of the starting *Fab-7^attP^* platform (Fig. 2C), this is not true for *5’P+scs^min^+HS3+PAS* (Fig. 3B). When the P-element promoter is included in the replacement, the A6 segment is almost completely absent: the A6 sternite is missing and there is only a rudimentary A6 tergite.

### Transcription also disrupts the functioning of the *iab-6* and *iab-5* regulatory domains

While the *scs^min^* replacement on its own has only minimal blocking activity, there were also some variable and unexpected LOF defects in the development of A5 (Fig. 2C). One plausible explanation for these LOF phenotypes is that transcription from the residual part of the *Cad87A* promoter disrupts the functioning of the *iab-5* domain as is observed for *iab-7* when *scs* is inserted in the reverse orientation.

To explore this possibility, we inserted *scs* in the forward orientation. We generated three different insertions, *scs* alone, *scs* plus HS3 (*scs+HS3*) and *scs* plus the three major *Fab-7* hypersensitive sites, HS1, HS2 and HS3 (*F7^HS1+2+3^*) (Fig. 4A). The *scs+HS3* and *scs+F7^HS1+2+3^* combinations have blocking activity and rescue the GOF transformations evident in the starting *F7^attP50^* platform. However, in both replacements A6 and A5 have an A4 like phenotype. This is most clearly seen in the pattern of pigmentation and in the dense trichome hairs in the A5 and A6 tergite (compare A5 and A6 with A4 in Fig. 4B). Consistent with the idea that transcriptional readthrough from the *Cad87* promoter interferes with the functioning of the *iab-5* and *iab-6* domains. RT-PCR experiments show that there are elevated levels of transcripts derived from *scs* in the *iab-5* and *iab-6* regulatory domains in the *scs+HS3* adult 2-days males compared to the *wt* 2-days males (Fig. S1).

**Fig. 4.**
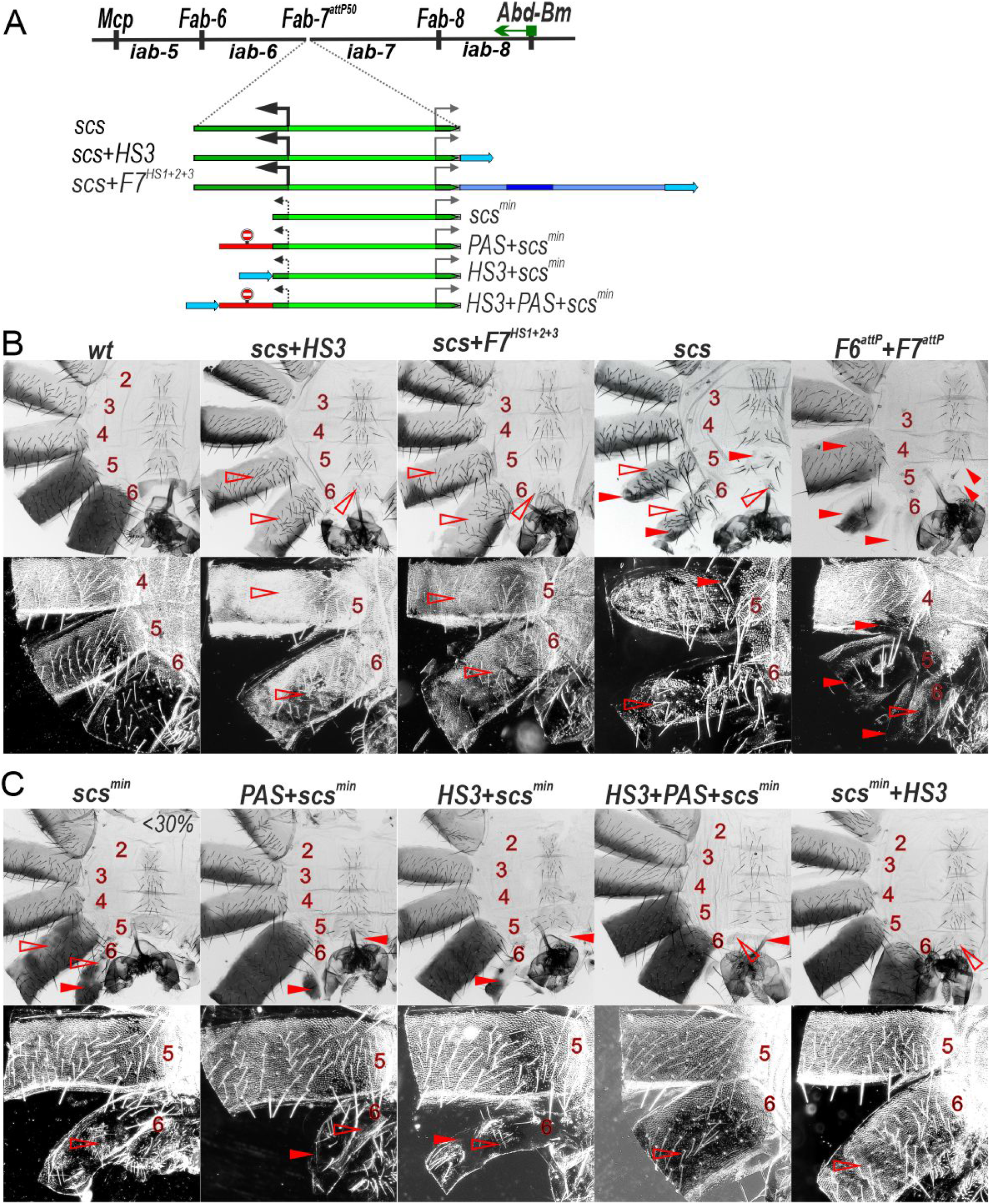
The *Cad87A* promoter in the *scs* inserted in the direct orientation (*scs*) is responsible for inactivation of the *iab-5* and *iab-6* domains. (A) *Fab-7* boundary replacement schemes in which *scs* and *scs^min^* were inserted in direct orientation. (B) Bright field (top) and dark field (bottom) images of cuticles prepared from males of *wt*, *scs, scs+HS3, scs+F7, F6^attP^+F7^attP^* Other designations are as in Fig. 2.

A more complicated phenotype is observed with *scs* alone (Fig. 4B). As expected, blocking activity is not complete and A6 shows evidence of GOF transformations. The A6 tergite is reduced in size, while there is only a small patch of sternite tissue. In both cases, the residual A6 tissue has a phenotype indicative of a transformation towards A4 identity: there are bristles on the patch of sternite tissue, while the residual tergite is depigmented and has patches of ectopic trichome hairs. Interestingly a mixed GOF and LOF phenotype is also observed in A5: both the sternite and tergite are reduced size as expected for a GOF transformation, while the tergite is depigmented and there are patches of densely packed trichome hairs. The GOF transformations in the *scs* replacement resemble those seen when both *Fab-7* and *Fab-6* are deleted (see *F6^attP^+F7^attP^* in Fig. 4B); however, unlike *scs* the double boundary deletion shows no evidence of LOF transformations of A5 and A6. There is also evidence of a weak GOF transformation of A4 in the double boundary deletion.

These findings indicate that transcription from the *Cad87* promoter in *scs* directed towards *iab-6* and *iab-5* disrupts the functioning of these domains. They also raise the possibility that the variable LOF phenotypes in A5 evident in males carrying the *scs^min^* replacement might be due a low level of transcription from the truncated *Cad87A* promoter. Indeed, in the immediate vicinity of the integration site (*attP/attB* fusion site), weak but verified transcripts from the *Cad87* promoter are found in *scs^min^* (Fig. S1). *scs^min^* does, however, differ from *scs* in that the level of transcripts in *scs^min^* (and *scs^min^+HS3*) in *iab-5* and *iab-6* is similar to background (Fig 1S). Since the LOF phenotypes in A5 varied between individuals and were seen in only about 30% of the *scs^min^* males, one plausible explanation is that stochastic differences in promoter activity between individuals might account for the incomplete penetrance. To test this possibility, we generated a *PAS+scs^min^* replacement (Fig. 4C). Unlike *scs^min^*, the A5 tergite in *PAS+scs^min^* males is fully pigmented in all adult males which would suggest that transcription from the clipped *Cad87* promoter is likely responsible for the pigmentation defects in A5. However, this does not seem to be true for the trichome hairs on the tergite as they are still densely packed like those in A4. This finding indicates that the trichome hair phenotype is likely due to the blocking activity of the *scs^min^* element, which prevents *iab-5* from regulating *Abd-B* in cells that can give rise to trichome hairs.

To further investigate the effects transcriptional readthrough, we placed HS3 upstream of *scs^min^* in *HS3+scs^min^*. Unlike *scs^min^+HS3, HS3+scs^min^* is unable to prevent crosstalk between *iab-6* and *iab-7* and A6 is transformed towards A7 (compare *scs^min^+HS3* with *HS3+scs^min^* in Fig. 4B). However, *HS3* is able to complement *scs^min^* when transcriptional readthrough is blocked by an interposed *PAS* sequence (*HS3+PAS+scs^min^*, Fig. 4B). Thus, a low level of transcription from the truncated *Cad87A* promoter is apparently sufficient to impact the boundary activity of *HS3*.

### Readthrough transcription disrupts the functioning of a minimal *Fub* replacement boundary

We wondered whether transcriptional readthrough would also impact the functioning of other boundary elements. To investigate this possibility, we chose the BX-C *Fub* boundary. *Fub* marks the border between the *Ubx* regulatory domain *bxd/pbx* and the *abd-A* gene and its regulatory domain, *iab-2* [32]. As illustrated in Fig. 5A, there are two *Fub* hypersensitive regions, HS1 and HS2. The larger *Fub* hypersensitive region HS2 contains motifs for several known chromosomal architectural proteins. The distal 177 bp HS2 sequence (*dHS1*) has binding sites for dCTCF and Su(Hw) and we found that it can function as an effective boundary [49,50]. The proximal 450 bp HS2 sequence (*pHS2*, Fig. 4A) contains binding sites for Pita and Su(Hw).

**Fig. 5.**
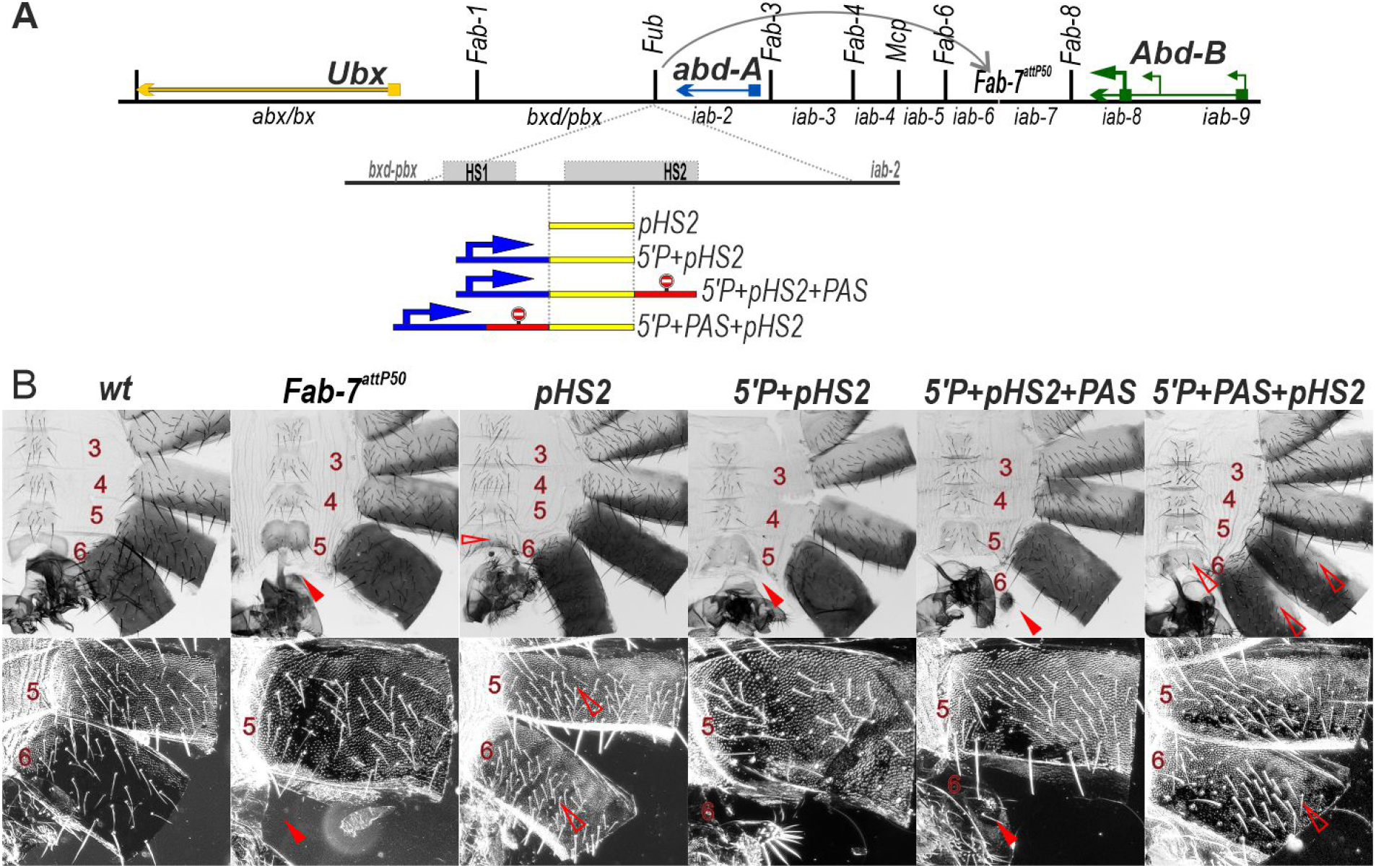
Transcription from the P-element promoter suppresses the activity of the *Fub* sub-fragment *pHS2*, when it replaces the *Fab-7* boundary. (A) *Fab-7* boundary replacement schemes, in which *pHS2* was inserted in different combinations with the P-element promoter and polyadenylation signal from SV40 (PAS). (B) Bright field (top) and dark field (bottom) images of cuticles prepared from males of *wt, Fab-7^attP50^, pHS2, 5’P+pHS2+PAS, 5’P+PAS+pHS2*. Other designations are as in Fig. 2.

We first tested whether *pHS2* is able to function as a boundary when introduced into the *Fab-7^attP50^* platform. As shown in Fig. 5B, *Fub pHS2* rescues the GOF phenotype of the *Fab-7^attP50^* deletion. Like most other heterologous replacements, *pHS2* blocks crosstalk but does not support bypass: an A6 segment is present in the *pHS2* replacement; however, its morphological features indicate that it has an A5 rather than an A6 identity. While the A5 tergite is fully pigmented, the trichome hairs are densely packed much like the A4 tergite (consistent with the idea that trichome morphology in A5 is more sensitive to blocking activity by replacement boundaries than pigmentation).

A different result is obtained when the P-element promoter is placed upstream of *pHS2* in the *Fab-7* replacement (Fig. 5B). As was observed for the P-element combination *5’P+scs^min^+HS3, pHS2* boundary activity is lost in *5’P+pHS2* and the A6 segment is missing. To test whether this is due to transcriptional readthrough from the P-element promoter we generated two additional replacement combinations. In the first the *PAS* element was placed downstream of the *pHS2* boundary to give *5’P+pHS2+PAS*, while in the second the *PAS* element was placed between the P-element promoter and *pHS2* to give *5’P+PAS+pHS2*. As would be expected if readthrough disrupts boundary function, there is only a residual A6 tergite in the *5’P+pHS2+PAS* replacement, while this GOF transformation is rescued when the *PAS* element is placed between the P-element promoter and *pHS2 (5’P+PAS+pHS2)* (Fig. 5B).

### Readthrough transcription disrupts *pHS2* function in its endogenous context

Bender and Fizgerald (2002) generated a series of imprecise hopouts of a P-element transgene inserted near the distal end of the *bxd/pbx* regulatory domain close to the sequences that were subsequently found to correspond to the *Fub* boundary [32,51]. These hopout events induced an anterior to posterior transformation of A1 towards A2 identity. Molecular characterization of one the hopouts that had a particularly strong phenotype, *Uab^HH1^*, revealed that it was a truncated P-element transgene that retained only the P-element promoter and 65 bp of *lacZ* coding sequence. The P-element transgene was also inverted so that the promoter was pointing towards the *Fub* boundary and the *abd-A* gene. Several potential mechanisms were proposed to account for the transformation of A1 to A2 induced by P-element transcription [51]. One was that transcription disrupted the functioning of an as yet unidentified boundary that blocked crosstalk between the *Ubx bxd/pbx* and *abd-A iab-2* regulatory domains. A second was that transcription interfered with the functioning of an element in *iab-2* that is required to keep the *iab-2* domain silenced in A1.

To test the boundary model, we took advantage of a *Fub* replacement platform *F2^attP^* (Fig. 6A) that removes a 2106 bp sequence containing the two nuclease hypersensitive sites associated with the *Fub* boundary and replacing it with an *attP* site (*F2^attP^*) [13]. As shown in Fig. 6B the A1 tergite in *wt* is narrower than the A2 tergite, lacks bristles and has less pigmentation, while the A1 sternite is absent. In *F2^attP^* males, the A1 segment is transformed into a copy of A2: the tergite is larger and it has a pigmentation and bristle pattern like A2, and there is also a sternite that is covered in bristles. These phenotypic transformations in the adult cuticle resemble those reported previously [51] for the P-element hopout mutants.

**Fig. 6.**
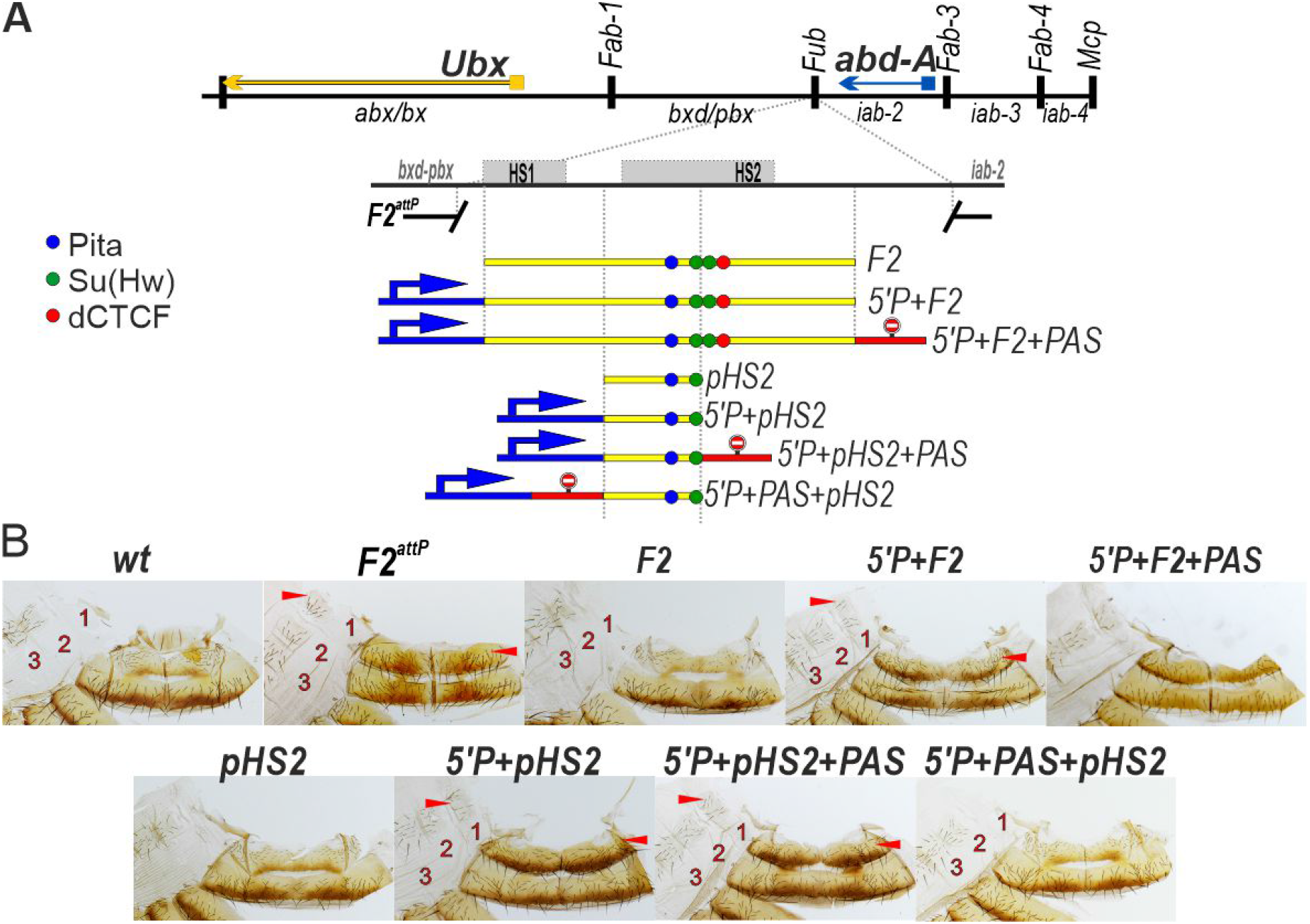
Transcription from the P-element promoter suppresses the functional activity of the *Fub* boundary and the *Fub* sub-element *pHS2* in its endogenous location between the *bxd/pbx* (the *Ubx* regulatory region) and *iab-2* (the *abd-A* regulatory region) domains. (A) *Fub* boundary replacement schemes, in which *Fub* or *pHS2* were inserted in different combinations with the P-element promoter and polyadenylation signal from SV40 (PAS). (B) Morphology of the male abdominal segments (numbered) in *wild type* (*wt*), *F2^attP^, F2, P+ F2, 5’P+F2+PAS* (C) Morphology of the male abdominal segments (numbered) in *wild type* (*wt*), *pHS2, 5’P+pHS2, 5’P+pHS2+PAS, 5’P+PAS+pHS2*.

The GOF transformations evident in the starting *F2^attP^* deletion platform can be fully rescued by a 1587 bp fragment, *F2*, which includes both HS1 and HS2 (Fig. 6A). Consistent the previous results [51], we find that rescuing activity is disrupted when the P-element promoter, *5’P* is placed upstream of the *F2* fragment (*5’P+F2*). In this replacement the A1 segment resembles A2 just like the starting deletion platform (Fig. 6B). Since introducing the PAS element downstream of *F2* in the *5’P+F2+PAS* combination does not rescue the GOF transformation, it would appear that boundary function rather than a downstream silencing element is the critical target for transcription inactivation.

To confirm these findings, we tested the *Fub pHS2* fragment used in the *Fab-7* replacement experiments. We found that *pHS2* on its own is sufficient to rescue the GOF transformations induced by the *F2^attP^* deletion – the A1 tergite is narrow, lacks bristles and is unpigmented while there is no A1 sternite as in *wt*. Rescuing activity is lost when the P-element promoter is placed upstream of *pHS2* in both *5’P+HS2* and *5’+pHS2+PAS*, and in both cases A1 is transformed towards an A2 identity. As would be expected if *pHS2* boundary function is disrupted by transcriptional readthrough from the P-element promoter, blocking activity is restored when the *PAS* element is placed between the P-element promoter and *pH2* (Fig. 6B).

## Discussion

### Blocking activity of scs is context dependent

Our results indicate that the *scs* boundary has only a limited ability to block crosstalk between the *iab-6* and *iab-7* regulatory domains. This result is unexpected, as in transgene assays *scs* was found to have one of the “stronger” insulator activities [42–44,52,53]. It seems likely that *scs* is a poor match with the neighboring *Fab-6* and *Fab-8* boundaries [54,55]. Both depend upon CTCF, while *scs* does not [56]. Also, when placed in the context of BX-C *scs* seems to have a cell and/or an enhancer specific blocking activity. For example, phenotype of the A5 tergite in *PAS+scs^min^* males (Fig. 4C) suggest that *scs* is unable to block the regulatory interactions between *iab-5* and *Abd-B* required for *wt* pigmentation, while its insulating activity is sufficient to block the interactions needed to inhibit the formation of trichomes.

### Transcription disrupts enhancer activity

In previous *Fab-7* replacement experiments [40], it was found that when *scs* is inserted in the reverse orientation, transcription from the *Cad87A* promoter towards *iab-7* induced a GOF transformation of A6(PS11) into A7(PS12). To explain this result, it was suggested that transcription through *iab-7* prematurely activated the domain. However, we found that when *scs^R^* was introduced into a larger *Fab-7* deletion that lacks HS3, transcription from the *Cad87A* interferes with the functioning of the *iab-7* domain, inducing a LOF transformation. That the activity of tissue specific enhancers in this region of BX-C is disrupted by transcriptional readthrough is supported by the effects of inserting *scs* in the direct orientation. In this case, it induces a LOF transformations of both A6(PS11) and A5(PS10) towards A4(PS9). The effects of transcription from the *Cad87A* promoter on these regulatory domains are most clear-cut when *scs* is combined with *HS3* or *F7^HS1+2+3^*. In both of these replacements the combination of *scs* with *HS3* or *F7^HS1+2+3^* suppresses the GOF phenotype of the *Fab-7^attP50^* deletion platform making the LOF transformations in A6(PS11) and A5(PS10) more obvious.

### Transcription disrupts boundary function

The enhancers in the *Abd-B* regulatory domains are not the only elements whose function is disrupted by transcriptional readthrough. We find that boundary activity can also be abrogated by readthrough transcription. In the case of our *Fab-7* replacements this is most directly demonstrated when boundaries are placed downstream of a P-element promoter. The *scs^min^+HS3* combination not only rescues the GOF transformation of A6(PS11) in the *Fab-7^attP50^* deletion platform, but also prevents *iab-6* and to a lesser extent *iab-5* domain from regulating *Abd-B* expression. However, if the P-element promoter is placed upstream of *scs+HS3* as in the *5’P+scs+HS3+PAS* combination, blocking activity is largely lost and A6 is transformed towards an A7 identity.

The effects of transcription on boundary activity are not limited to *scs* and *HS3 (Fab-7*) as transcription also interferes with the functioning of the *Fub* boundary fragment *pHS2*. On its own it rescues the GOF phenotype of *Fab-7^attP50^*; however, when placed downstream of the P-element promoter, blocking activity is lost. Transcription also inactivates the *Fub* boundary in its native context. This is true for both the 412 bp *pHS2* element, and a larger 1587 bp *Fub* fragment (Fig. 6). Moreover, as was case for a combination in which the truncated *Cad87A* promoter in *scs^min^* is pointing towards *HS3*, the disruption in *pHS2* boundary function by the P-element promoter can be rescued by placing the *PAS* element in between the promoter and the boundary. These findings argued that boundary function is disrupted by readthrough transcription rather than some other properties of the promoter. Similar results are observed when *pHS2* or a larger *Fub* fragment were tested for rescuing the *Fub* deletion. Again, the rescuing activity is lost when *pHS2* and *F2* are combined with the P-element promoter. Thus, the effects of transcription also do not appear to be context dependent.

While previous studies have shown that readthrough transcription can suppress the activity of enhancers and promoters, how this happens is not fully understood [57–59]. One idea is that RNA Pol II transiently displaces DNA binding proteins as it passes [60,61]. In the case of boundary elements, it seems possible that even a transient displacement of factors important for their activity could have a significant impact on boundary function. Fly boundaries link distant sequences together to form looped domains or TADs by boundary:boundary pairing interactions [5,11,14]. In this mechanism, TADs are formed when proteins associated with one boundary element physically interact in a stable fashion with proteins associated with a second boundary element. This means that a transient displacement of boundary associated proteins from one of the elements would disrupt the TAD as it would uncouple the physical linkage between the distant sequences that define endpoints of the loop. Consistent with this idea, a low level of transcription from the truncated *Cad87A* promoter can disrupt the boundary functions of *HS3*.

While our experimental paradigm is artificial, there are contexts in which transcriptional readthrough provides a mechanism for coordinating higher order chromosome organization with regulating gene activity. For example, the blocking activity of the *Fub-1* boundary is turned off by readthrough of a lncRNA from a promoter that is activated by the *Ubx* regulatory domain *bxd/pbx* [62] in PS6/A1 and more posterior parasegments. Inactivation of the *Fub-1* boundary enables enhancers in the *bxd/pbx* domain to regulate *Ubx* expression. MicroC experiments suggest that transcriptional readthrough of the *Fub-1* boundary is likely accompanied by a switch from one TAD configuration to another configuration. Since transcription is not continuous, but instead occurs in bursts that can differ both in their length and frequency depending on the specific enhancer: promoter combinations, a readthrough mechanism would result in only a transient remodeling of the TAD organization. Moreover, this remodeling would also be subject to regulation. In this respect it is interesting to note that lncRNAs are thought to account for a vast majority of the transcripts in mammalian genomes [63–66]. It would be reasonable to suppose that some of these lncRNAs span TAD boundaries (as well as other elements like “tethering” elements that might also be sensitive to readthrough). Transcriptional readthrough of these lncRNAs would then alter the local TAD organization and in doing so generate new combinations of regulatory elements and potential target genes.

## Materials and methods

### Generation of the replacement lines

The strategy of the *Fab-7* replacement lines is described in detail in [48,67]. The *F2^attP^* replacement is described in detail in [13]. DNA fragments used for the replacement experiments were generated by PCR amplification and verified by sequencing. The sequences of the used fragments are shown in the Supporting Table S1.

### Cuticle preparations

Adult abdominal cuticles of homozygous eclosed 3±4 day old flies were prepared essentially as described in [39]. Photographs in the bright or dark field were taken on the Nikon SMZ18 stereomicroscope using Nikon DS-Ri2 digital camera, processed with ImageJ 1.50c4 and Fiji bundle 2.0.0-rc-46.

### RNA purification and quantitative analysis

For each replicate, 20 adult 2- to 3-day-old males were collected and frozen in liquid nitrogen. Total RNA was isolated using the TRI reagent (MRC) according to the manufacturer’s instructions. RNA was treated with DNAse I (Thermo Scientific) to eliminate residual genomic DNA. The synthesis of cDNA was performed using RevertAid Reverse Transcriptase (Thermo Scientific) in a reaction mixture containing 5 μg of RNA and 5 μM random hexamer. The amounts of specific cDNA fragments were quantified by real-time PCR. At least three independent biological replicates were made for each experiment. At least four independent technical replicates were made for each RNA sample. Relative levels of mRNA expression were calculated in the linear amplification range by calibration to a standard curve. RNA levels were normalized to a level of housekeeping gene Vha100-1. The sequences of oligonucleotides used in the study are presented in Table S2.

## Acknowledgments

We thank Farhod Hasanov for fly injections.

## Author Contributions

**Conceptualization:** Pavel Georgiev, Olga Kyrchanova

**Data curation:** Olga Kyrchanova

**Funding acquisition:** Paul Schedl, Olga Kyrchanova

**Investigation:** Olga Kyrchanova, Vladimir Sokolov, Maxim Tikhonov

**Methodology:** Olga Kyrchanova, Maxim Tikhonov

**Project administration:** Paul Schedl, Olga Kyrchanova, Pavel Georgiev

**Resources:** Paul Schedl, Olga Kyrchanova

**Supervision:** Paul Schedl, Pavel Georgiev, Olga Kyrchanova

**Validation:** Olga Kyrchanova, Paul Schedl, Pavel Georgiev

**Visualization:** Olga Kyrchanova, Vladimir Sokolov, Maxim Tikhonov

**Writing original draft:** Paul Schedl, Olga Kyrchanova, Pavel Georgiev

**Writing review & editing:** Paul Schedl, Olga Kyrchanova, Pavel Georgiev

## Supporting information captions

**S1 Fig.**
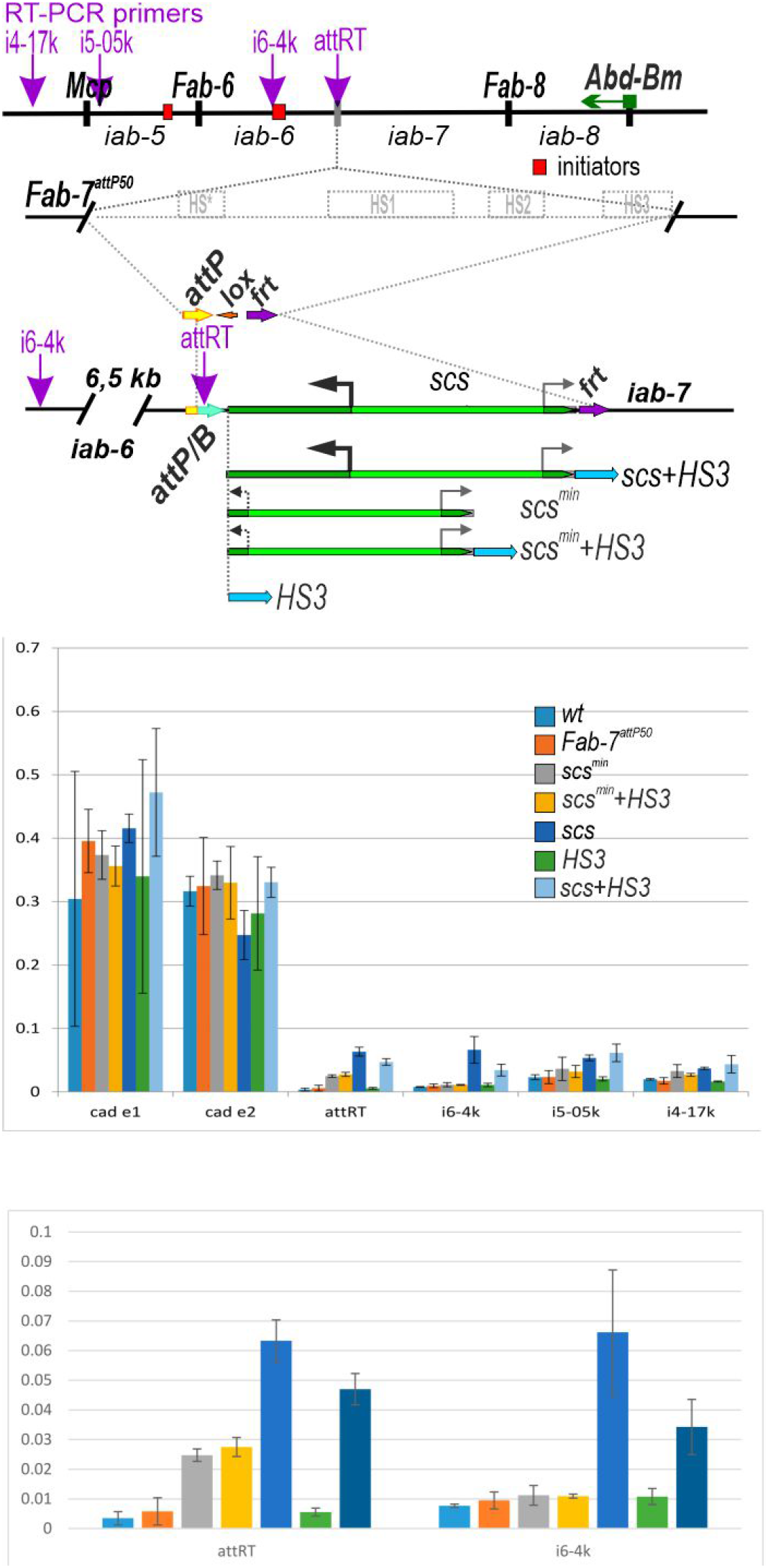
Detection transcription from *Cad87A* promoter in *Fab-7* replacements with *scs* and *scs^min^*. The results of real-time PCR (RT-PCR) show the level of detected transcripts (in both orientations) that are detected at different testing sites as indicated (shown with magenta arrows on *Abd-B* regulatory region scheme). Points “Cad e1” and “Cad e2” are taken from *Cad87A* 1^st^ exon and reflect transcription from genomic and transgene promoters. The results are presented as a percentage of input genomic DNA. Error bars show standard deviations of triplicate PCR measurements for three independent experiments.

**S1 Table.**
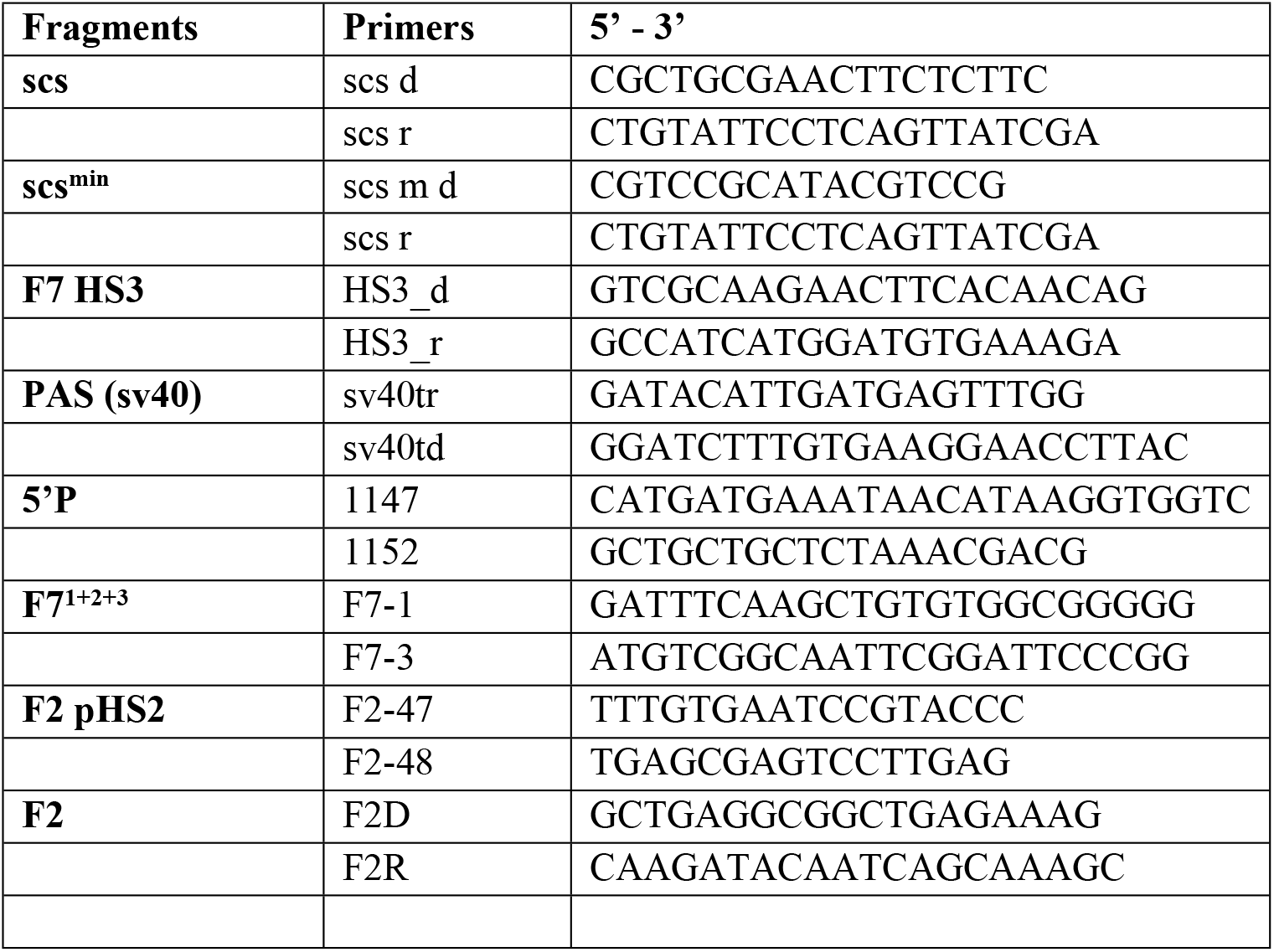
Primers for generating fragments.

**S2 Table.**
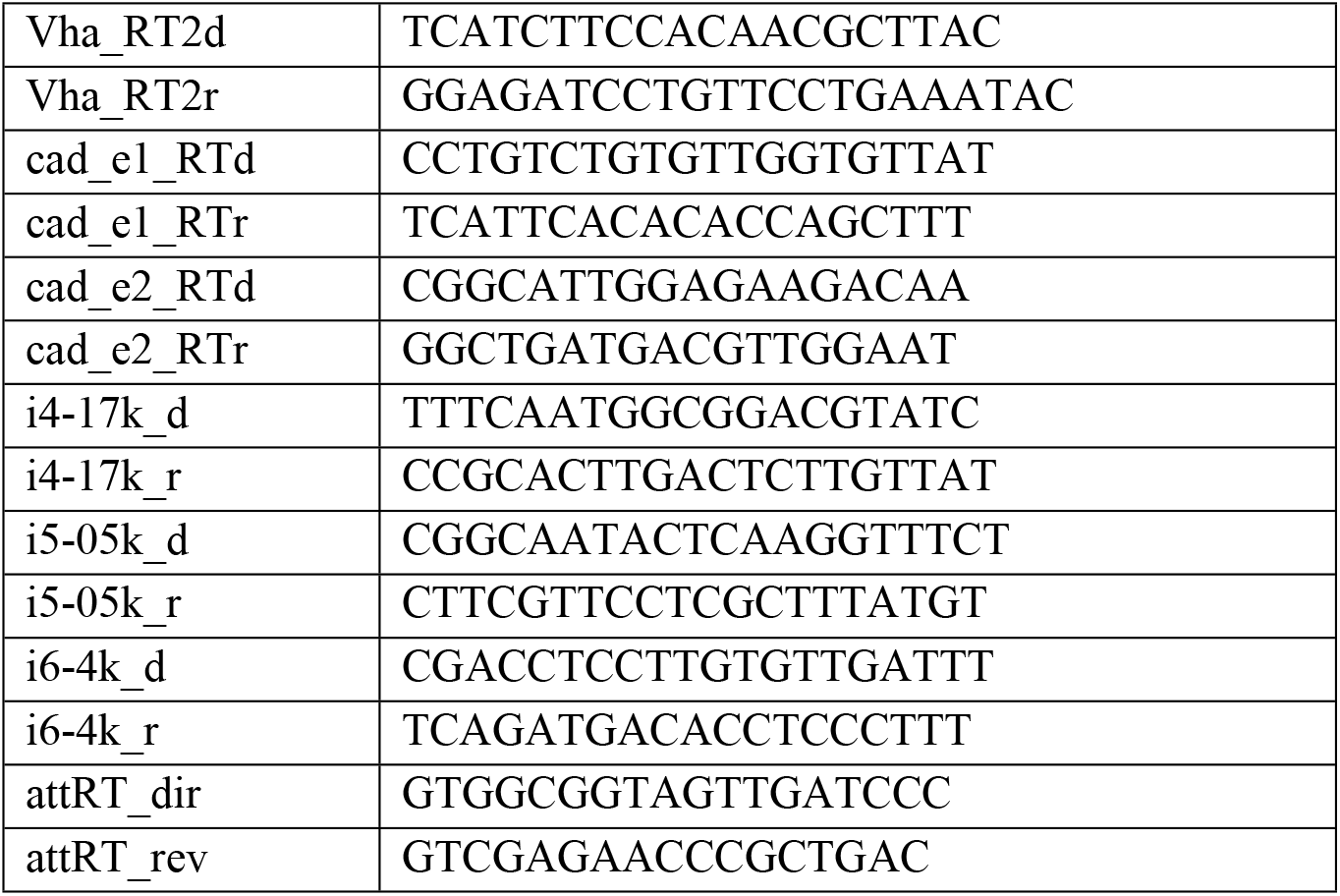
The sequences of oligonucleotides used in real-time PCR.

